# Phosphorylation at Ser65 modulates ubiquitin conformational dynamics

**DOI:** 10.1101/2022.08.09.502833

**Authors:** Remy A. Yovanno, Alvin Yu, Tyler J. Wied, Albert Y. Lau

**Affiliations:** Department of Biophysics and Biophysical Chemistry, Johns Hopkins University School of Medicine, 725 N. Wolfe Street, WBSB 706, Baltimore, MD 21205, USA; Department of Chemistry, The University of Chicago, 5735 S. Ellis Avenue, Chicago, IL 60637, USA

**Keywords:** ubiquitin, phosphorylation, molecular dynamics, string method

## Abstract

Phosphorylation of ubiquitin at Ser65 increases the population of a rare C-terminally retracted (CR) conformation. Transition between the CR and the Major ubiquitin conformations is critical for promoting mitochondrial degradation. The mechanisms by which the Major and CR conformations of Ser65-phosphorylated (pSer65) ubiquitin interconvert, however, have not yet been revealed. Here, we perform all-atom molecular dynamics simulations using the string method with swarms of trajectories to calculate the lowest free-energy path between these two conformers. Our analysis reveals the existence of a Bent intermediate in which the C-terminal residues of the β5 strand shift to resemble the CR conformation, while pSer65 retains contacts resembling the Major conformation. This stable intermediate was reproduced in well-tempered metadynamics calculations, with the exception of a Gln2Ala mutant that disrupts contacts with pSer65. Lastly, dynamical network modelling reveals that the transition from the Major to CR conformations involves a decoupling of residues near pSer65 from the adjacent β1 strand.

## INTRODUCTION

Ubiquitin (Ub) phosphorylation at Ser65 by protein kinase 1 (PINK1) initiates the mitophagy pathway through activation of the E3 ligase Parkin^1–6^. PINK1 aggregates on the cytosolic surface of depolarized mitochondria^7^ where it phosphorylates ubiquitin^1^. The phosphorylated ubiquitin (pUb) then binds to and activates Parkin^4–6^, which mediates the assembly of polyubiquitin chains on mitochondrial outer membrane proteins^8^, recruiting autophagy receptors and ultimately forming the LC3-positive phagophore for mitochondrial degradation^9–11^. Dysfunction of PINK1 and Parkin are associated with early-onset autosomal-recessive Parkinson’s disease^11–13^. In addition, Ser65 pUb granules have been found in post-mortem human brain samples that are increased by age and with Parkinson’s disease^14,15^.

Although the first crystal structure of ubiquitin was published in 1985^16^, recent NMR studies have revealed a new conformation of ubiquitin in which the C-terminal beta strand, *β*5, retracts by two amino acids^17,18^. Both ubiquitin and Ser65- phosphorylated (pSer65) ubiquitin exist in an equilibrium between this C-terminally retracted (CR) conformation and the Major (Maj) conformation captured by the original crystal structure. The two conformations primarily differ by the network of hydrogen bonds formed between strand *β*5 and adjacent strands, *β*1 and *β*3^17^. Chemical exchange saturation transfer (CEST) NMR experiments report that in phosphorylated ubiquitin, the occupancy of the Major (pUb-Maj) and CR (pUb-CR) conformations are 70% and 30%, respectively, whereas in non-phosphorylated ubiquitin, the population of the CR conformation (Ub-CR) is only ~0.68%^18^. This indicates that phosphorylation at Ser65 modulates the conformational dynamics of ubiquitin. Transition between these two conformations is critical for initiating mitophagy; PINK1 phosphorylates ubiquitin at Ser65 in the CR conformation^18^, while the Major conformation of pUb is required to bind to and activate Parkin^6,18^. Although previous work has identified a transition pathway between the Ub-Maj and Ub-CR conformations^19^, little is known about the molecular mechanism driving the transition between the pUb-Maj and pUb-CR conformations. Understanding how pUb conformational dynamics differ from Ub at a molecular level would provide valuable insight into the unique role of pUb in pathogenesis.

Here, we compute the transition path between pUb-Maj and pUb-CR and identify important intermediates along the pathway. We then compute conformational free energy landscapes for both pUb and Ub to probe how Ser65 phosphorylation affects the ubiquitin conformers that are sampled. Lastly, we explore how the extent of local dynamic coupling involving the residues of *β*5 plays an important role in driving the overall conformational transition.

## RESULTS AND DISCUSSION

### Transition pathway for pUb

To study the transition between the pUb-Maj and pUb-CR conformations, we computed the transition mechanism using all-atom molecular dynamics simulations employing the string method with swarms of trajectories^20–22^. Starting from the pUb-Maj conformation, the phosphate group of pSer65 interacts with Gln62 **(Fig. S1A)**. pSer65 then flips outward toward the *β*1 strand. Hydrogen bonds between pSer65 and Gln62 break, and the protonated phosphate contacts the Lys63 backbone carbonyl **(Fig. S1B)** and occasionally the polar atoms of Gln2 **(Fig S1C)**. The pSer65 phosphate remains flipped toward the *β*1 strand when the *β*5 strand shift is initiated **(Fig 1A)**.

**Fig. 1.**
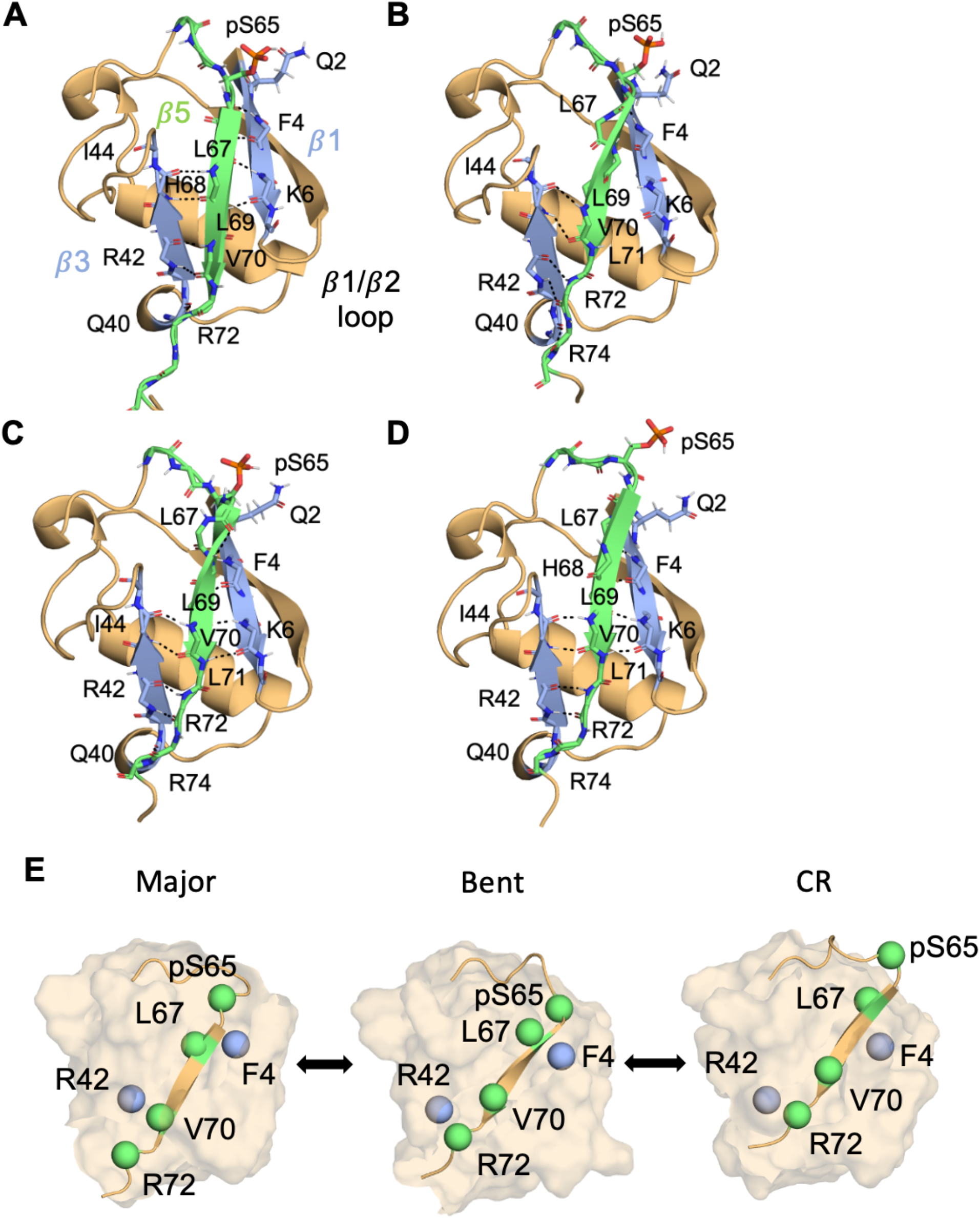
Conformational transition between the Major and CR conformations of pUb. **(A)** Prior to the *β*5 strand shift, pUb exists in the Major conformation where the pSer65 phosphate is flipped toward the *β*1 strand. **(B)** pUb adopts a Bent conformation by partial strand shift breaking interactions between *β*5 residues (His68, Val70, and Arg72) and *β*3 residues (Gln40, Arg42, and Ile44). This shift results in new interactions formed between *β*5 residues (Val70, Arg72, and Arg74) and *β*3 residues (Gln40, Arg42, and Ile44). Leu69 shifts toward Leu67, and contacts between those residues and the *β*5 strand are broken. Retention of the hydrogen bond between the pSer65 carbonyl oxygen and the Phe4 backbone amine contributes to strand strain. **(C)** New contacts form between Lys6 and both Leu69 and Leu71. The Phe4 backbone carbonyl forms a new contact with the Leu69 backbone amine. *β*5 strand tension is relieved when the hydrogen bond between the pSer65 backbone carbonyl and the Phe4 backbone amine is broken. **(D)** pUb adopts the CR conformation by forming a new hydrogen bond between the Leu67 backbone carbonyl and the backbone amine of Phe4. pSer65 swings away from Gln2. **(E)** Summary cartoon highlighting the key features of the pUb transition mechanism.

The *β*5 strand shift mechanism involves the following changes **(Fig 1B)**. (1) Interactions involving residues 68-72 (*β*5) with residues 40-44 (*β*3) shift to resemble those that are characteristic of the retracted conformation. However, the *β*5 residues N-terminal to His68 remain mostly unshifted, causing the strand to begin to twist/bend. (2) The Leu67 backbone carbonyl breaks interactions with the backbone amine of Lys6, and the Leu67 backbone amine breaks interactions with the backbone carbonyl of Phe4. However, the Phe4 backbone amine remains bound to the backbone carbonyl of pSer65. (3) Leu69 shifts toward Leu67. When Leu69 moves, its backbone amine breaks contact with the carbonyl of Lys6.

In the next step, new interactions form between the backbone carbonyl of Leu69 and the backbone amine of Lys6; the carbonyl of Lys6 is free to form new interactions with the Leu71 backbone amine. The newly free Phe4 backbone carbonyl now forms interactions with the Leu69 backbone amine, which causes the remaining hydrogen bond with the Phe4 backbone amine and pSer65 carbonyl to break **(Fig. 1C)**. In this *β*5 Bent state, the pSer65 phosphate oxygen can interact with the Gln2 sidechain amide.

The Phe4 backbone amine forms interactions with the Leu67 backbone carbonyl. pSer65 then swings away from Gln2, leaving it totally solvent exposed, resulting in the retracted conformation **(Fig. 1D)**. The overall transition between pUb-Maj and pUb-CR can be likened to the movement of an inchworm; the *β*5 strand first shifts from its C-terminal end causing it to bend and then from its N-terminal end, which relieves strand tension **(Fig. 1E)**.

We then compared the pUb transition mechanism to that previously reported for Ub^19^. Both transitions are initiated by the breaking of interactions at the C-terminal end of the *β*5 strand. However, the pUb mechanism involves an initial rotation of pSer65 away from Gln62 before any other *β*5 strand rearrangements occur. In the Ub mechanism, the initiation of this transition results from shifts in the *β*1/*β*2 loop that allow Leu71 to approach Leu69 and initiate the strand shift. The *β*5 strand then breaks all interactions between the *β*1 and *β*3 strands before shifting to the Ub-CR conformation. Unique to the pUb mechanism is the Bent intermediate in which the hydrogen bonding pattern between *β*5 residues 68-72 and *β*3 residues 40-44 resembles the pUb-CR conformation, while backbone interactions between pSer65 and Phe4 from the pUb-Maj conformation are maintained. We hypothesize that this intermediate is stabilized by interactions between the pSer65 phosphate group and the Gln2 sidechain.

Since the string method is ideal for capturing local structural change but may not fully sample global transitions, we next probed the energetics of these conformers using metadynamics free-energy calculations to validate the mechanisms.

### Energetics of pUb and Ub conformational transitions

Previous work comparing the structure and dynamics of pUb and Ub have indicated that pUb occupies the CR conformation much more readily than Ub^18^. This difference in Major and CR populations between pUb and Ub indicate different energetics associated with the transition. However, little is known about the molecular determinants of this difference. To probe the conformational thermodynamics of this transition, we computed free energy landscapes for both pUb and Ub using well-tempered metadynamics^23^. For these calculations, we employed a two-dimensional order parameter, (*q*_1_, *q*_2_), that captures the transition observed in our pUb pathway **(Fig. 2A,B)**. The first, *q*_1_, was used in previous work to describe the *β*5 shift relative to the *β*1 strand^19^ (see Methods). A small *q*_1_, value indicates the Major conformation, and a large *q*_1_, value indicates the CR conformation. However, our pUb mechanism indicates that the shift of the C-terminal half of *β*5 shifts relative to the *β*3 strand before the hydrogen bonding network with *β*1 rearranges to resemble the CR conformation. To account for this in our metadynamics simulations, we introduced an additional order parameter, *q*_2_, that captures this mechanistic detail by monitoring the distance between the C-terminal residues of *β*5 (Val70 and Arg72) and the *β*3 strand. A small *q*_2_ value is consistent with the Major conformation, and a large *q*_2_ value is consistent with both the Bent and CR conformation. The resulting free energy landscapes for pUb **(Fig. 2C)** and Ub **(Fig. 2D)** support the thermodynamic significance of the Bent intermediate. In our pUb energy landscape, we observe extensive sampling and increased stability of the Bent intermediate. In contrast, our Ub energy landscape shows a small population of conformers in which the pSer65 bends in the opposite direction (away from the *β*1 strand) without C-terminal retraction. According to the exchange rate computed from CEST NMR experiments^18^, the energy barrier between the Major and CR conformations is 19.32 kcal mol^-1^ (at 45°C) for Ub and 17.83 kcal mol^-1^ for pUb (at 25°C) assuming one-step Arrhenius behavior. These values are within the ranges captured by our computed energy landscapes.

**Fig. 2.**
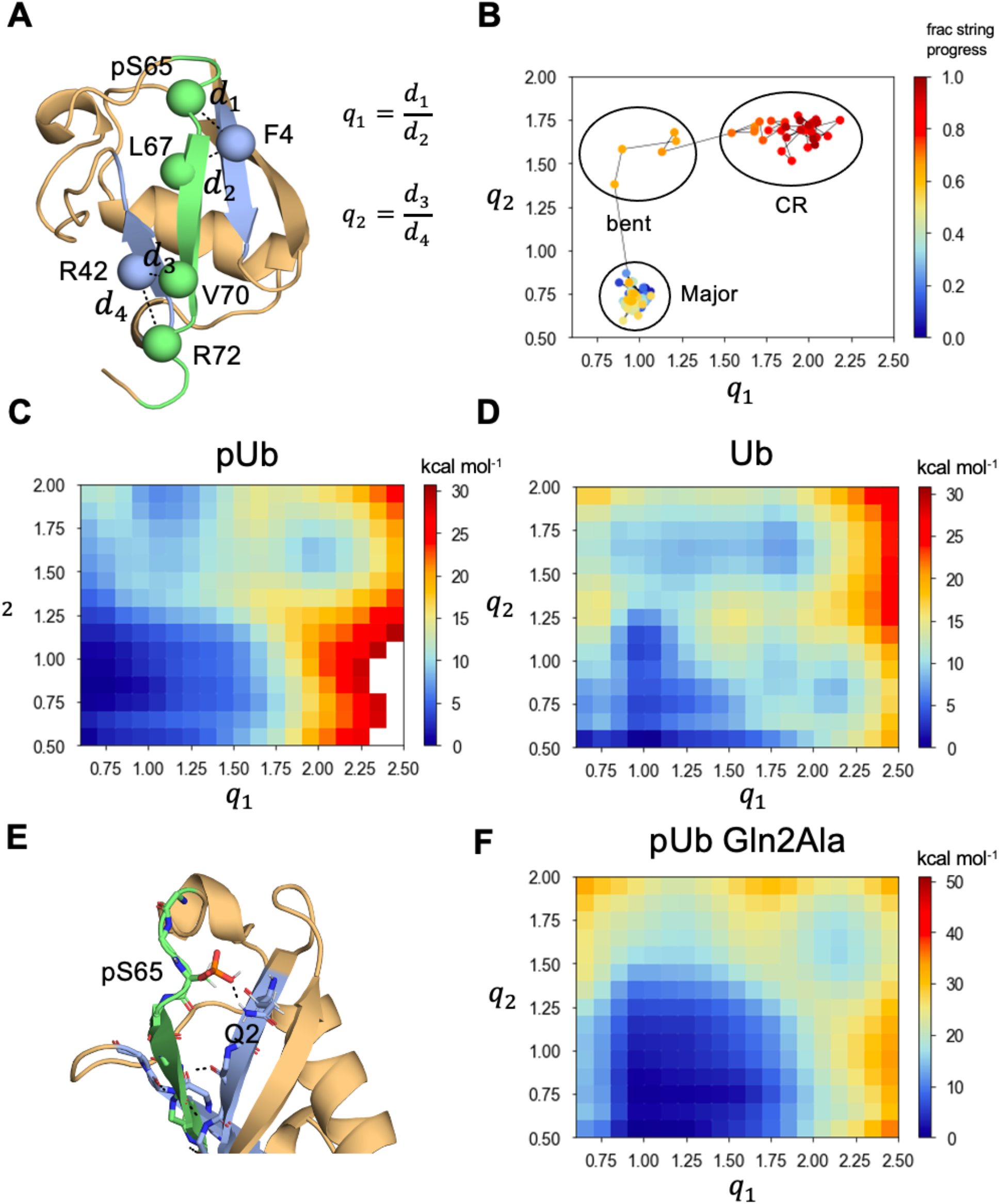
Free energy landscapes for pUb and Ub. **(A)**Visual representation of the two-dimensional order parameter (*q*_1_, *q*_2_) used for metadynamics simulations. **(B)** Steps of the pUb transition mechanism (computed from the string method) mapped onto the (*q*_1_, *q*_2_) order parameter to illustrate how this order parameter captures the structural changes observed throughout the transition. The color bar monitors the string progress where dark blue is the initial pUb-Maj structure and red is the final pUb-CR structure. **(C)**Free energy landscape (potential of mean force) computed for pUb. **(D)** Free energy landscape (potential of mean force) computed for Ub. **(E)** Hydrogen bond between the phosphate group of pSer65 and the Gln2 sidechain. **(F)** Free energy landscape (potential of mean force) computed for the pUb Gln2Ala mutant.

The transition pathway computed between pUb-Maj and pUb-CR reveals a Bent intermediate that is characteristic of the pUb transition but not for that previously determined for Ub. The presence of the Bent intermediate in the energy landscapes supports our pUb mechanism obtained using the string method, suggesting that large, global removal and re-insertion of *β*5 from the beta sheet is not required for transition. According to our energy landscapes, the Bent conformation is more stable relative to the CR conformation for pUb than for Ub. On a molecular level, the Bent conformation of pUb features a hydrogen bond between the pSer65 phosphate group and the Gln2 sidechain that is not present in the absence of a phosphorylated Ser65 **(Fig. 2E).** A charged-polar hydrogen bond is about 2 kcal mol^-1^ stronger than a polar-polar hydrogen bond^24^, which can account for the difference between pUb and Ub. To probe whether this hydrogen bond stabilizes the pUb-Bent conformation, we performed a metadynamics simulation of a Gln2Ala pUb mutant and observed that the Bent conformation is significantly less stable, ~20 kcal mol^-1^ less than the pUb-Maj conformation **(Fig. 2F)**. The CR conformation also shows decreased stability relative to the pUb-Maj conformation. This indicates that the pSer65–Gln2 interaction is critical for stabilizing the pUb-Bent conformation and facilitating the Maj-to-CR transition.

### Dynamic coupling for different pUb conformations

We next explored the local dynamics of each of the important intermediates identified along the pUb transition pathway by performing 100 ns of equilibrium molecular dynamics simulations and applying a dynamical network model to quantify the strength of coupled interactions involving the residues of *β*5. Specifically, we performed simulations of both pUb and Ub starting from the (1) Major **(Fig. 3A,B)**, (2) Bent **(Fig. 3C,D)**, and (3) CR **(Fig. 3E,F)** conformations to compare the contacts formed with *β*5. For each contact pair, we computed a mutual-information-based correlation coefficient^25^ that quantifies the degree of coupling between the two residues. The dynamical network data support the presence of coupling between *β*5 and *β*1, and *β*5 and *β*3, involving both polar and hydrophobic residues. The transition from the Major conformation to the CR conformation involves the decoupling of either Ser65 or pSer65 ((p)Ser65) with *β*1 and *β*3. (p)Ser65 coupling is present in the Major conformation, decreased in the Bent conformation and absent from the CR conformation. The shifting of interactions at the C-terminal half of *β*5 causes (p)Ser65 to “break out” of the beta sheet. A similar weakening of coupled interactions occurs with Leu67. The strand shift also engages *β*5 residues Leu73, Arg74, and Gly75 in coupled interactions primarily with *β*3. In addition, pSer65 of pUb-Bent exhibits weak coupling with Phe4, which is absent from Ser65 of the Ub-Bent conformation. Combined with results from our metadynamics simulations, it appears that stronger pSer65 coupling to *β*1 due to the hydrogen bond with Gln2 stabilizes the Bent conformation; decoupling results when Gln2 rotates away from *β*5, facilitating the remainder of the transition to the CR state.

**Fig. 3.**
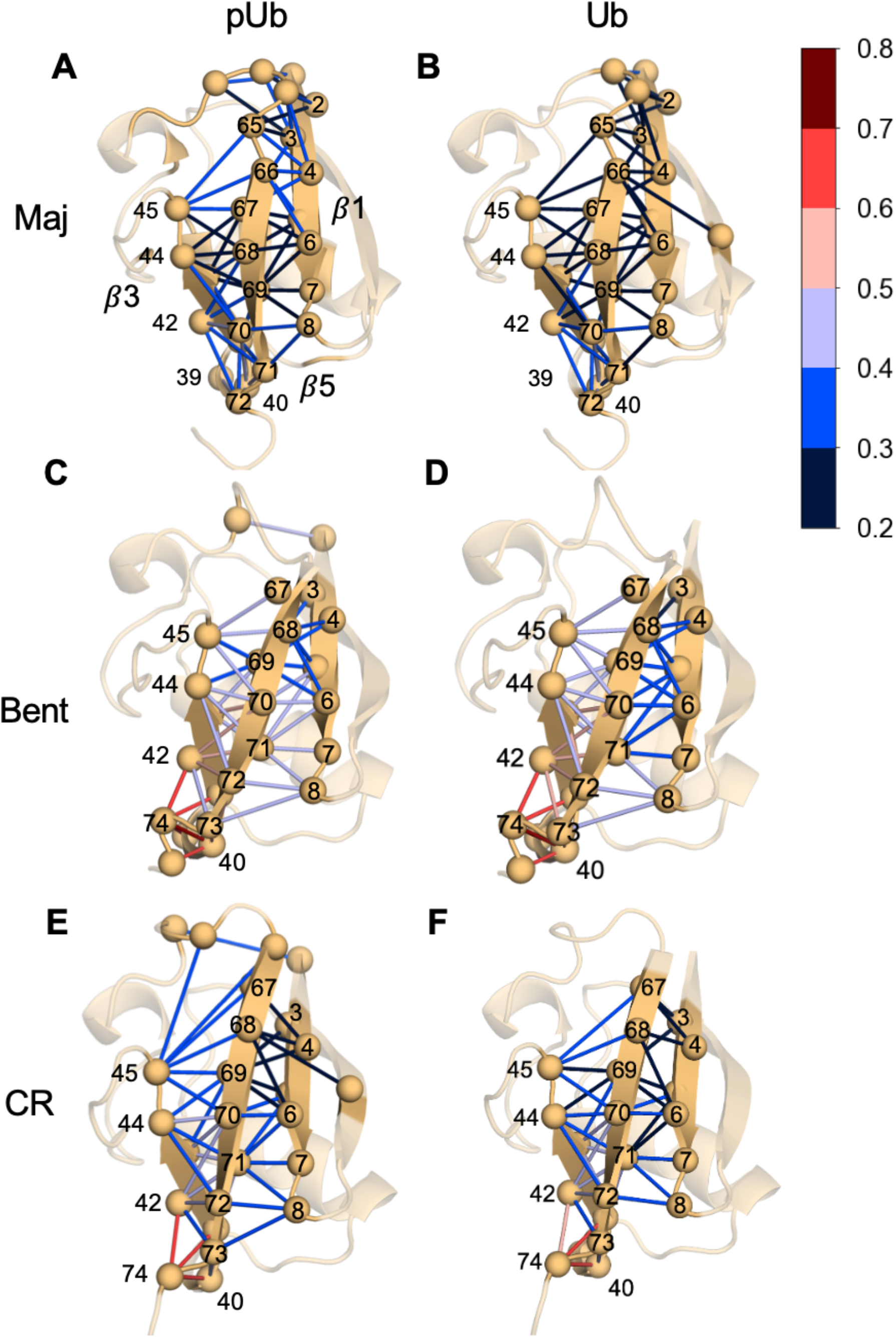
Dynamic coupling between residues of the *β*5 strand and the adjacent *β*1 and *β*3 strands. Dynamical network representation generated from equilibrium molecular dynamics simulations of important pUb and Ub conformational states. Value ranges of the generalized correlation coefficients are indicated by the colors of the edges connecting *C_α_* atoms in the network. A dynamical network was generated for **(A)** pUb in the Major conformation, **(B)** Ub in the Major conformation, **(C)** pUb in the Bent conformation, **(D)** Ub in the Bent conformation, **(E)** pUb in the CR conformation, and (**F)** Ub in the CR conformation. Correlograms illustrating these data are shown in **Figs. S6** and **S7**.

A prior simulation study with ubiquitin cites the disorder of the *β*1/*β*2 loop as the primary factor contributing to the initiation of the transition from Ub-Maj to Ub-CR^19^. We examined the extent of disorder of this loop for both pUb and Ub and computed the root mean square fluctuation (RMSF)^26^ for loop residues. We found that the RMSF for *β*1/*β*2 loop residues is higher than all other regions of the protein except for the *β*5 strand **(Fig. S8A)**. In our network data, we see that Leu8 is initially coupled most strongly to *β*5 residues Val70 and Ile71 in the Major conformation. Upon transition to the Bent conformation, we observe Leu8 coupling with both Arg72 and Leu73 that is also present in the CR conformation. This coupling between residues of the *β*5 strand and the *β*1/*β*2 loop are echoed in a previous molecular-dynamics simulation study of ubiquitin in the Major conformation that examined causal relationships between residue fluctuations using transfer entropy^27^. According to these data, solvent-exposed Glu64 at the N-terminal end of *β*5 transfers entropy to Ile3, which is accepted by an entropy sink, Leu8. The neighboring residue Thr7 of the *β*1/*β*2 loop drives the motion of *β*5 residue Leu71^27^. Our network analysis indicates that Glu64-Ile3 coupling is present in the Major conformation but is not present in the Bent or CR states of pUb or Ub. Combining the mutual-information-based couplings with these causal relationships, we find that information flow between the *β*5 and *β*1 strands is an important component for initiating C-terminal retraction and suggests that local dynamics help promote large-scale conformational motion.

## CONCLUSIONS

Here, we investigated the effects of Ser65 phosphorylation on the transition mechanism and conformational dynamics of ubiquitin. We applied the string method with swarms of trajectories to compute the minimum free energy path between the Major and CR conformations of pUb and discovered that the *β*5 strand “crawls” upward like an inchworm, first from its C-terminal residues and then breaking residues pSer65 and Leu67. This mechanism differs from a previously proposed Ub transition mechanism^19^, which involves shifts in a concerted fashion and does not feature a Bent intermediate. From our computed free energy landscapes, we determined that both pUb and Ub sample the Bent conformation. However, interactions between Gln2 and the phosphate group of pSer65 further stabilize the Bent state for pUb. Lastly, our network analysis of molecular dynamics trajectories indicates that the transition to the CR conformation involves the decoupling of residues at the N-terminal end of *β*5 and that allosteric communication between *β*5 and *β*1 is essential for initiating this transition from the Major to CR conformations of pUb and Ub. In total, this work provides insights into the molecular mechanisms driving the conformational dynamics of pUb.

## METHODS

### Model construction

All-atom models of pUb were constructed from crystal structures of the Major (PDB ID: 4WZP^17^) and CR (PDB ID: 5OXH^18^) conformations. Missing residues were added using the ModLoop Server^28^. Since the crystal structure of the CR conformation uses a TVLN mutant construct (T66V and L67N), these sidechains were converted back to their WT sequence using SCWRL4^29^.

### String method

The transition pathway for Ser65 pUb was computed using the string method with swarms of trajectories^21^, which uses short unbiased trajectories to determine the minimum free energy path (MFEP) between two stable states along a set of order parameters. An initial transition path between pUb-Maj and pUb-CR was generated using PYMOL’s Morph feature^30^, yielding an initial string of 100 images (conformers) including pUb-Maj (img_0) and pUb-CR (img_99). Coordinates of the *C_α_* atom for each residue were used as order parameters for computing the string. Each string image was solvated in an electrostatically neutral orthorhombic box of size 62 Å × 44 Å × 40 Å containing 150 mM NaCl and 10618 atoms in total. A weak center-of-mass harmonic restraint of 0.5 kcal mol^-1^ Å^-2^ was applied to the N, CA, and C atoms of residues 23, 26, and 30 to prevent large translational and rotational protein motion. The SP1 phosphorylation patch in the CHARMM27 force field was applied to Ser65 to generate the proper topology. A CHARMM^31^ implementation of the string method developed in previous work^22^ was used to perform the calculations. <u>All</u> simulations in this work were performed at 310 K using the CHARMM36 force field with the SP1 patch from CHARMM27.

Each cycle of the string method consisted of 5000 steps of restrained equilibration centered at the target value for each order parameter, 500 steps of restrained dynamics to generate the initial configuration from which to initiate the trajectory swarms, and 10 trajectories of 50 steps of unrestrained dynamics from which the average drift was computed to obtain the next iteration of the string. This resulting string was then re-parameterized to ensure images were equally distanced in collective variable space. String convergence was assessed by calculating the string RMSD (here, the *C_α_* RMSD) for each cycle using the initial string as the reference **(Fig. S2A)**. To correct for translational and rotational protein motion that contributes to the RMSD calculation, string RMSD was computed after all conformers were aligned to img_0, more clearly illustrating that the string has converged **(Fig. S2B)**. Hydrogen bond networks for each string image were computed using MDAnalysis^32,33^.

To ensure reproducibility of the key mechanistic steps, the string calculation was repeated with 34 images and 100 swarms. The converged string was able to reproduce the Bent intermediate **(Fig. S3)**.

### Metadynamics

Well-tempered metadynamics^23^ simulations were performed along two collective variables: *q*_1_ describes the ratio of distances between the *C_α_* atoms of Phe4 and (p)Ser65 to Phe4 and Leu67, and *q*_2_ describes the ratio of distances between Arg42 and Val70 to Arg42 and Arg72. These calculations were performed with Gaussian half-widths of 0.1 for each collective variable, an initial hill height of 0.2 kcal mol^-1^, and a bias factor of 10. The Gaussian biases were deposited every 2 ps. Metadynamics simulations for pUb, Ub, and the Gln2Ala pUb mutant were initiated from the Bent conformation to ensure this conformation was sampled. Each system was equilibrated in an NVT ensemble with gradually relaxing backbone and sidechain restraints and followed by a nanosecond of NPT equilibration prior to applying the metadynamics bias. A weak center-of-mass harmonic restraint of 0.5 kcal mol^-1^ Å^-2^ was applied to the N, CA, and C atoms of residues 23, 26, and 30. All metadynamics simulations were performed using GPU-accelerated NAMD^34^. Error in the energy landscapes was computed by calculating the average potential of mean force at each (*q*_1_, *q*_2_) at 50 ns increments throughout the last 500 ns of metadynamics simulations **(Fig. S4)**. The initial 3 *μs* metadynamics simulation of the Gln2Ala pUb mutant failed to fully sample the CR conformation; the results of this simulation were included in **Fig. S5**.

### Equilibrium molecular dynamics and dynamical network analysis

Equilibrium molecular dynamics were performed using the GPU-accelerated NAMD^34^ simulation package. Simulations were initiated from Major, Bent, and CR conformations for both pUb and Ub. To ensure the protein did not exceed the overall system extent, a solvent box of 70 Å × 70 Å × 70 Å was used for each simulation. Each system was electrostatically neutral and contained 150 mM NaCl and approximately 33,000 atoms. Equilibration was performed in an NVT ensemble with gradually relaxing backbone and sidechain restraints. Production simulations were carried out in an NPT ensemble at 1 atm. Trajectories were aligned by their backbone atoms, and frames were sampled at 10 ps intervals for analysis.

A dynamical network model was constructed from the equilibrium molecular dynamics trajectories using the dynetan python package^35^. To generate the network, a node is defined by each *C_α_* atom in the protein, and a pair of nodes is connected by an edge if the heavy atoms of both residues are within 4.5 Å of each other for at least 75% of the trajectory. The strength of each edge was determined by calculating a generalized correlation coefficient, *r_MI_*, as a function of a mutual information estimator, *I*, constructed by counting the number of frames in which the “distance” (maximum variation of the three dimensions) for each node is within a cutoff defined from the maximum variation of the *k* nearest neighbors to a reference frame that is varied throughout the trajectory^25^. The generalized correlation coefficient is then computed from *I* using the following equation for *d*=3 dimensions:

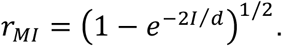

This approach to determining inter-node correlation is useful because it accounts for non-linear relationships in node fluctuations^36^. The 100 ns trajectory for each system was divided into four windows from which the standard error of each pairwise correlation value was reported **(Tables S1-S6)**.

## Acknowledgements

We used resources provided by the Maryland Advanced Research Computing Center (MARCC) and Advanced Research Computing at Hopkins (ARCH) at Johns Hopkins University. This work was funded by the Johns Hopkins Catalyst Award (to A.Y.L.); NIH T32GM135131 (to R.A.Y.).

## Author Contributions

R.A.Y., A.Y., T.J.W., and A.Y.L. designed the research. R.A.Y. conducted molecular dynamics simulations; R.A.Y. and A.Y.L. analyzed the results; R.A.Y., A.Y., and A.Y.L. wrote the manuscript.

## Competing Interests

The authors declare no competing interests.

## Notes

### Competing Interest Statement

The authors have declared no competing interest.

