## Supplemental Information for "Phosphorylation at Ser65 modulates ubiquitin conformational dynamics"


**Supplemental information includes:**

**Figures S1-S8**

**Tables S1-S6**

**Movie S1:** pUb Major to CR transition mechanism calculated by the string method with swarms of trajectories, related to Fig. 1.

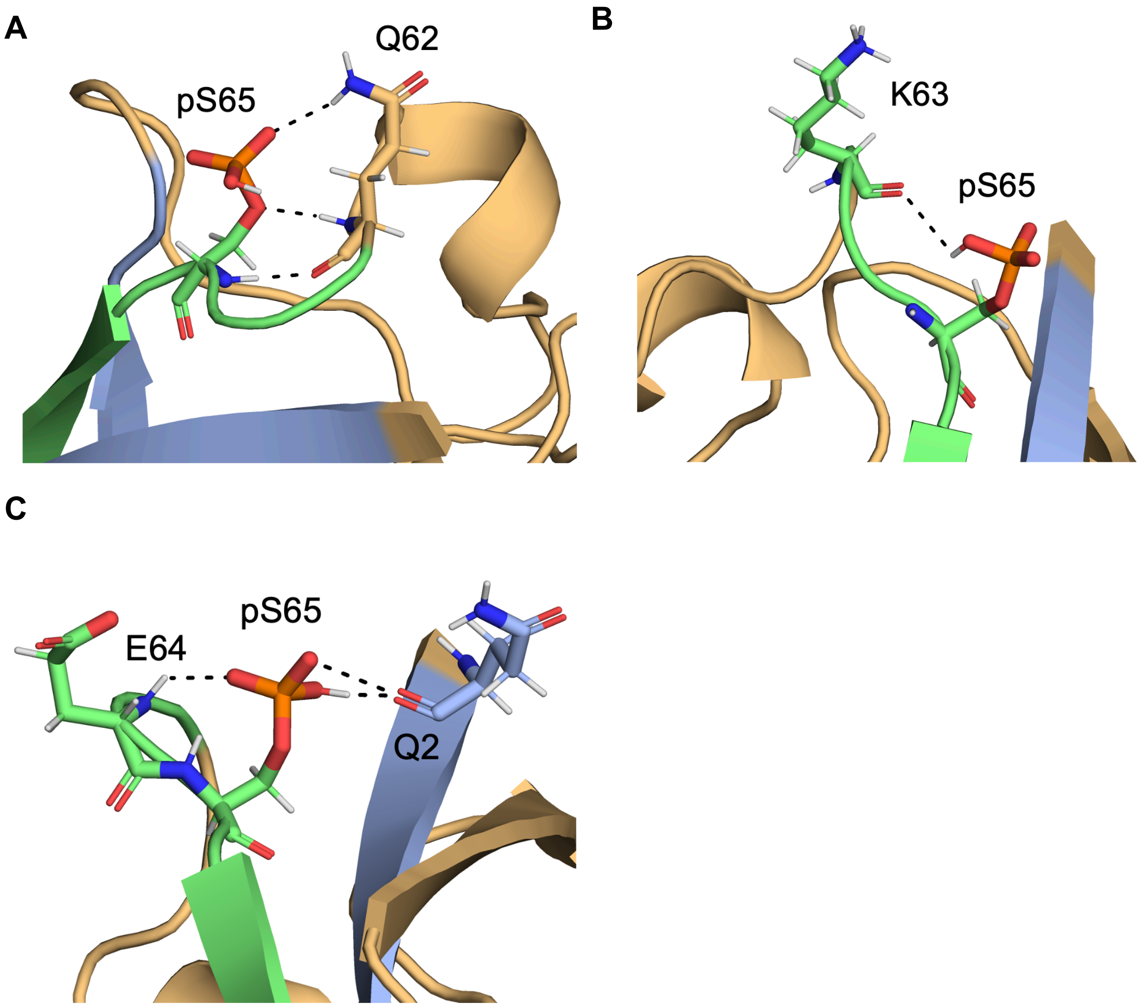


**Fig. S1. Analysis of the transition pathway computed by the string method, related to Fig. 1.** Local dynamics of the pUb Major conformation including **(A)** interaction between pSer65 and Gln62. **(B)** interaction between the pSer65 phosphate group and the Lys63 backbone carbonyl. **(C)** Interactions formed by the pSer65 phosphate with both Gln2 and Glu64 backbones.

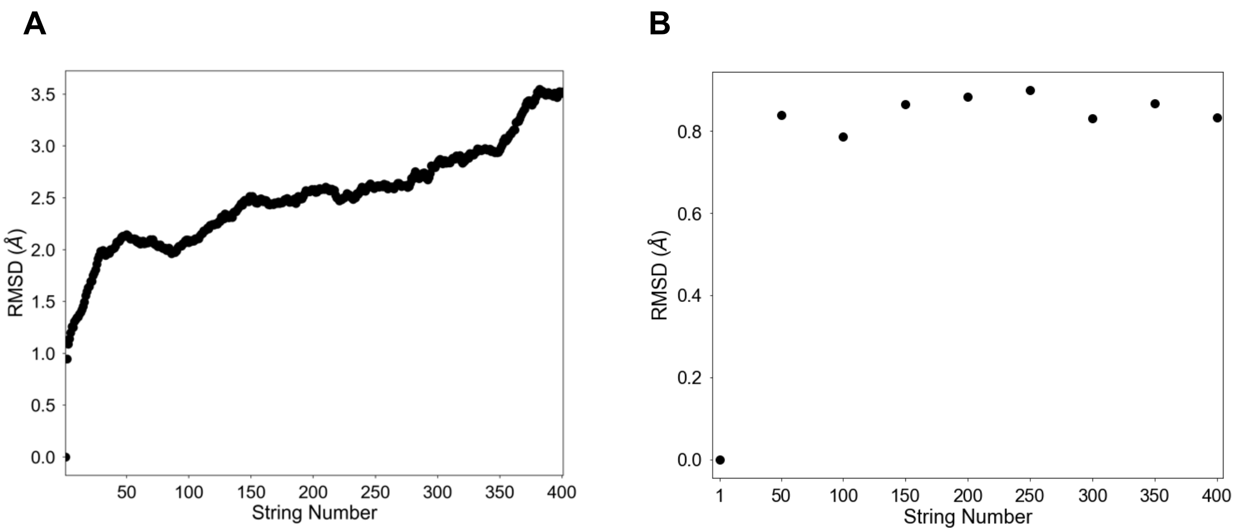


**Fig. S2. Assessing string convergence, related to Fig. 1. (A)** String RMSD for each cycle relative to the initial string used for assessing string convergence. **(B)** String RMSD computed after alignment to the backbone atoms of img_0 (Major) to correct for translational and rotational protein motion.

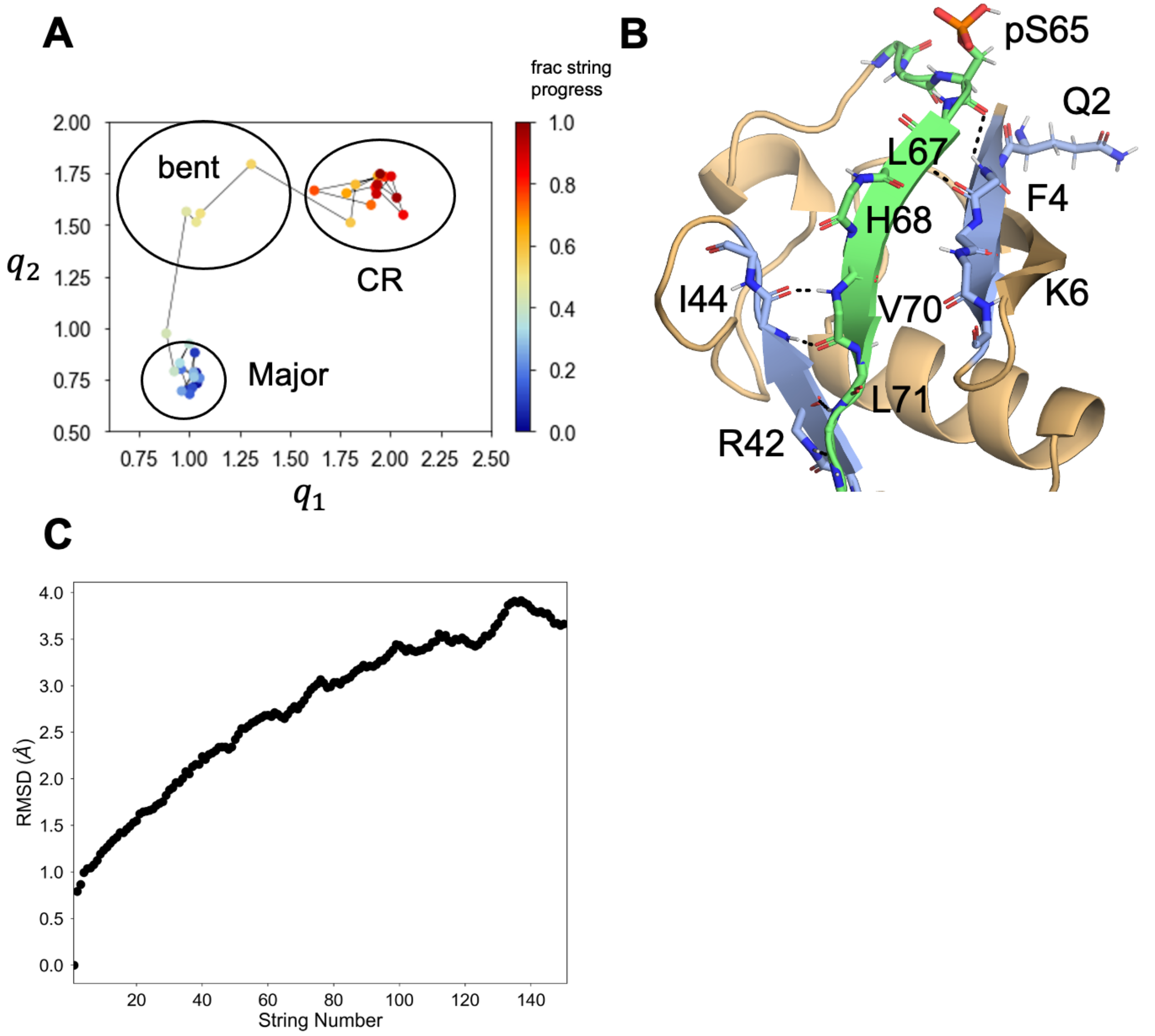


**Fig. S3. Converged string using 34 images and 100 swarms, related to Figs. 1 and 2. (A)** Our $(q_1, q_2)$ order parameter was computed along the converged string at cycle 150 to illustrate that the transition mechanism is reproduced with different parameters. **(B)** A conformer representing the bent state. **(C)** String RMSD to demonstrate convergence.

,

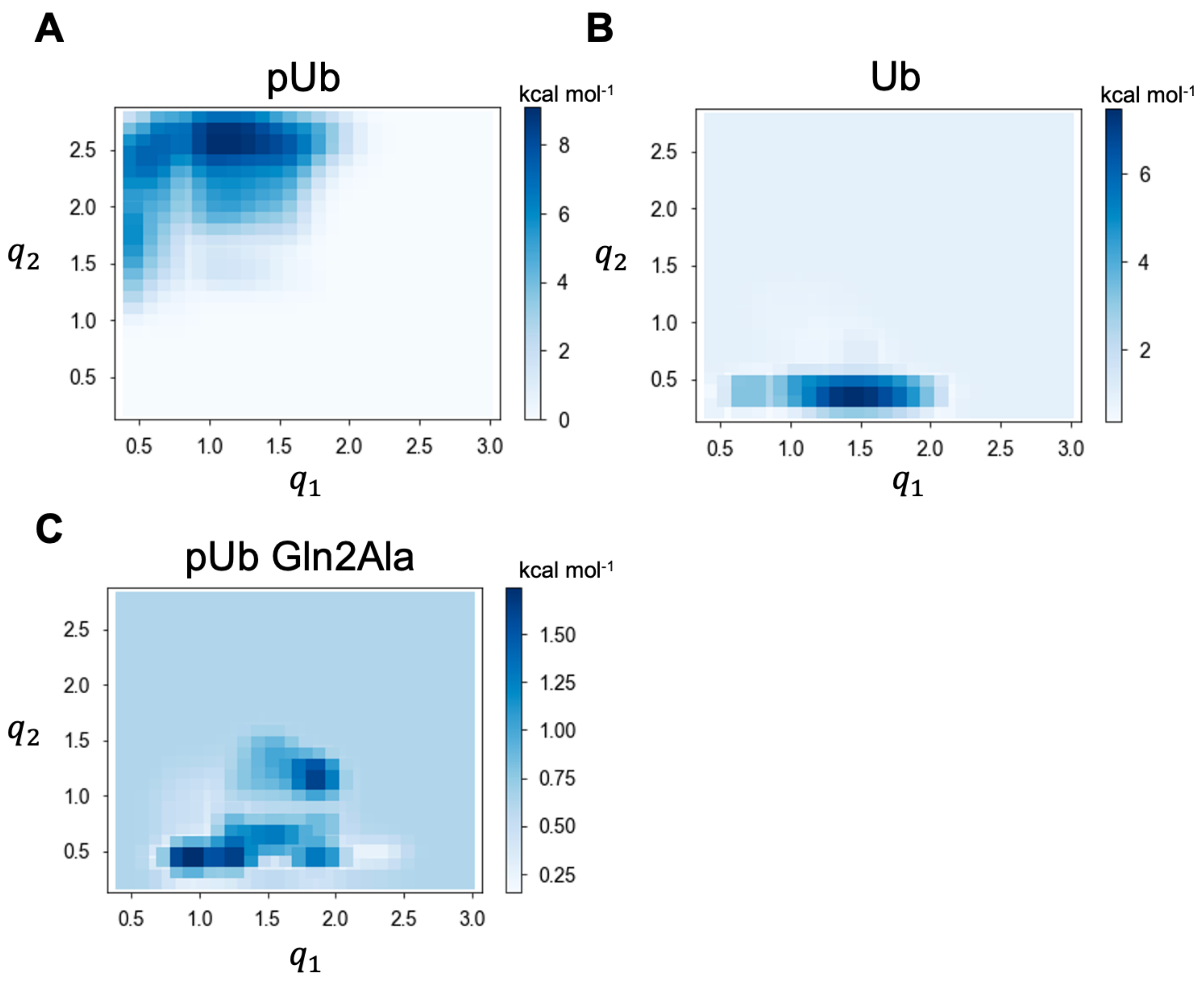


**Fig. S4. Metadynamics error plots, related to Fig. 2.** Standard deviations of the energy landscapes (potential of mean force) computed for each $(q_1, q_2)$ value over the last 500 ns at 50 ns intervals. Error plots were computed for **(A)** pUb, **(B)** Ub, and **(C)** the Gln2Ala pUb mutant. Regions of high standard deviation are outside of the regions defined by the relevant conformers sampled by the string outlined in **Fig. 2**.

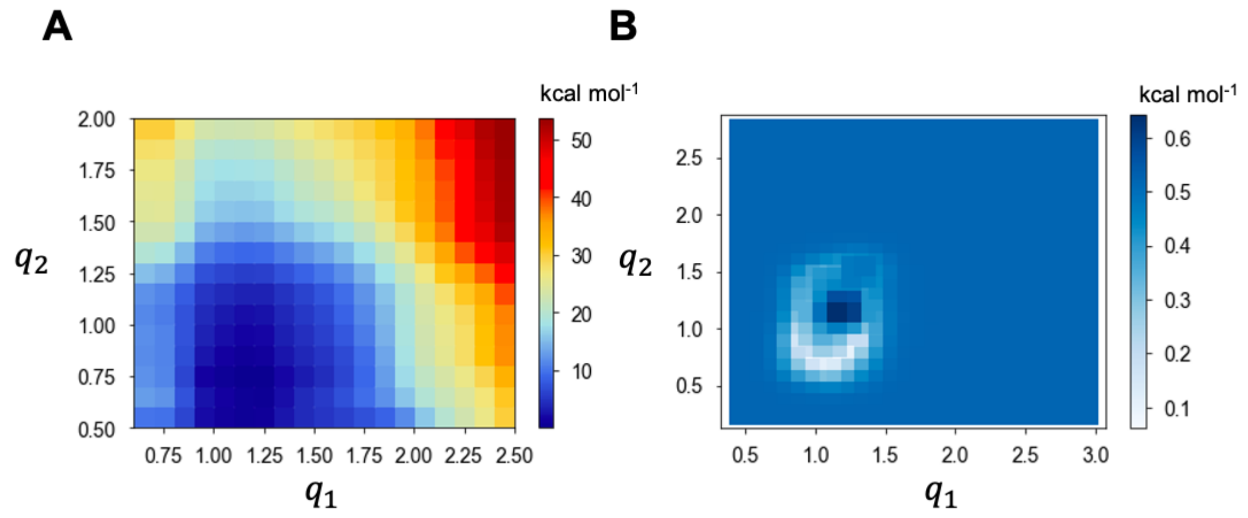


**Fig. S5. Partial conformational sampling of Gln2Ala-pUb from a metadynamics simulation, related to Fig. 2. (A)** PMF and **(B)** error plot for a preliminary metadynamics trial seems to have converged without fully sampling the CR state (upper right corner).

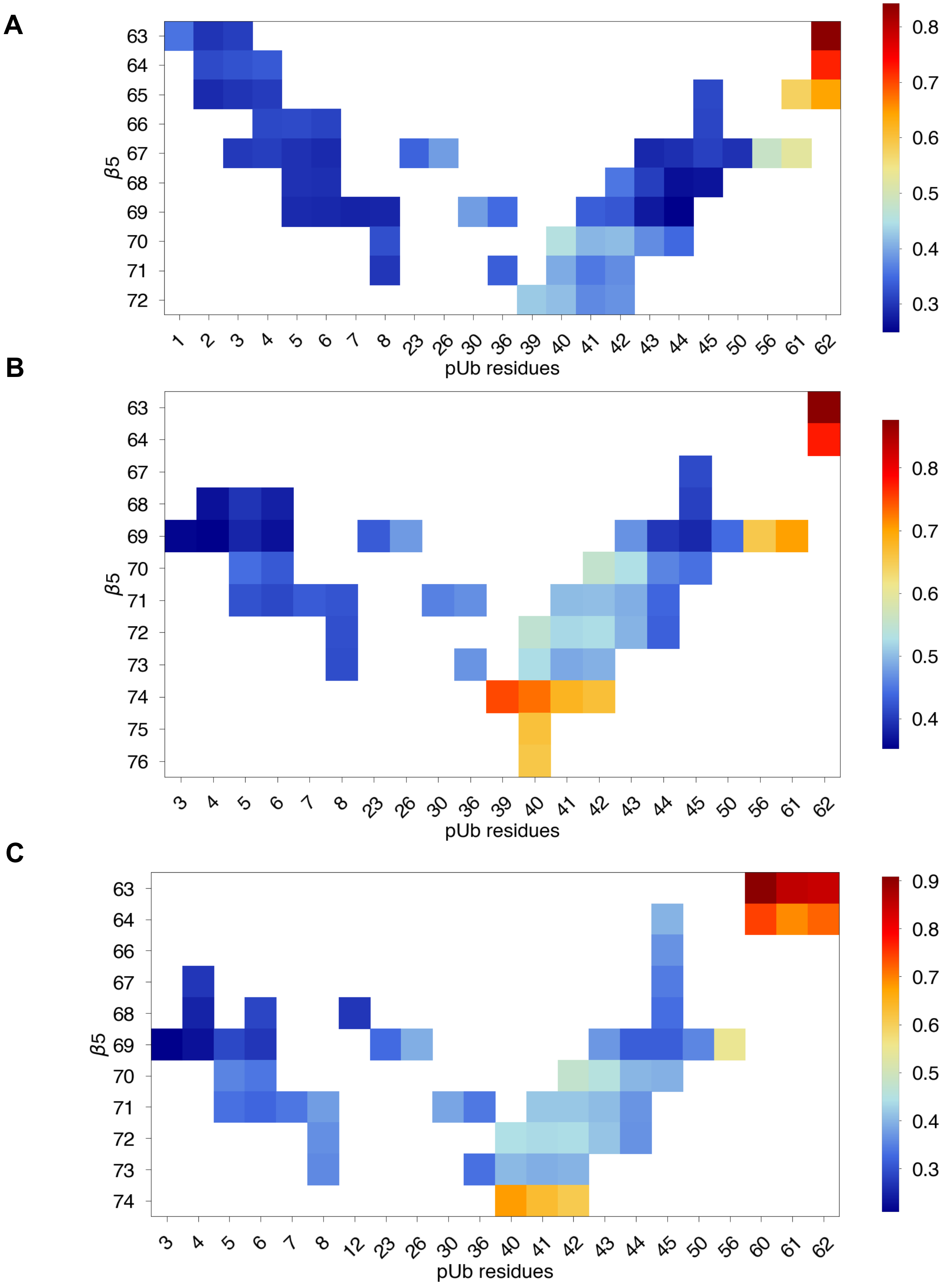


**Fig. S6. Correlograms for pUb residues interacting with the $\beta$5 strand, related to Fig. 3.** Correlation matrices are shown for **(A)** Major, **(B)** Bent, and **(C)** CR conformations.

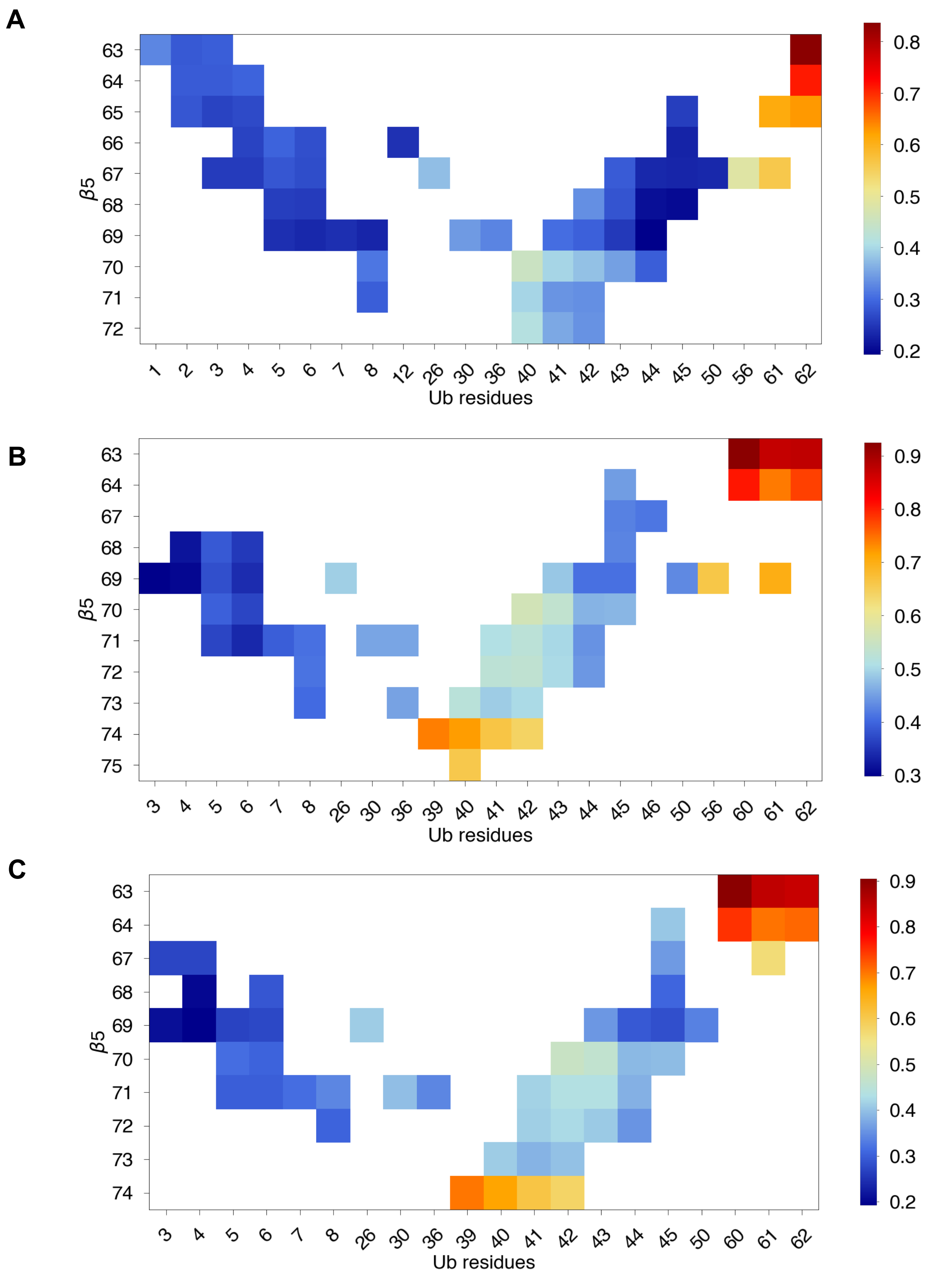


**Fig. S7. Correlograms for Ub residues interacting with the $\beta$5 strand, related to Fig. 3.** Correlation matrices are shown for **(A)** Major, **(B)** Bent, and **(C)** CR conformations.

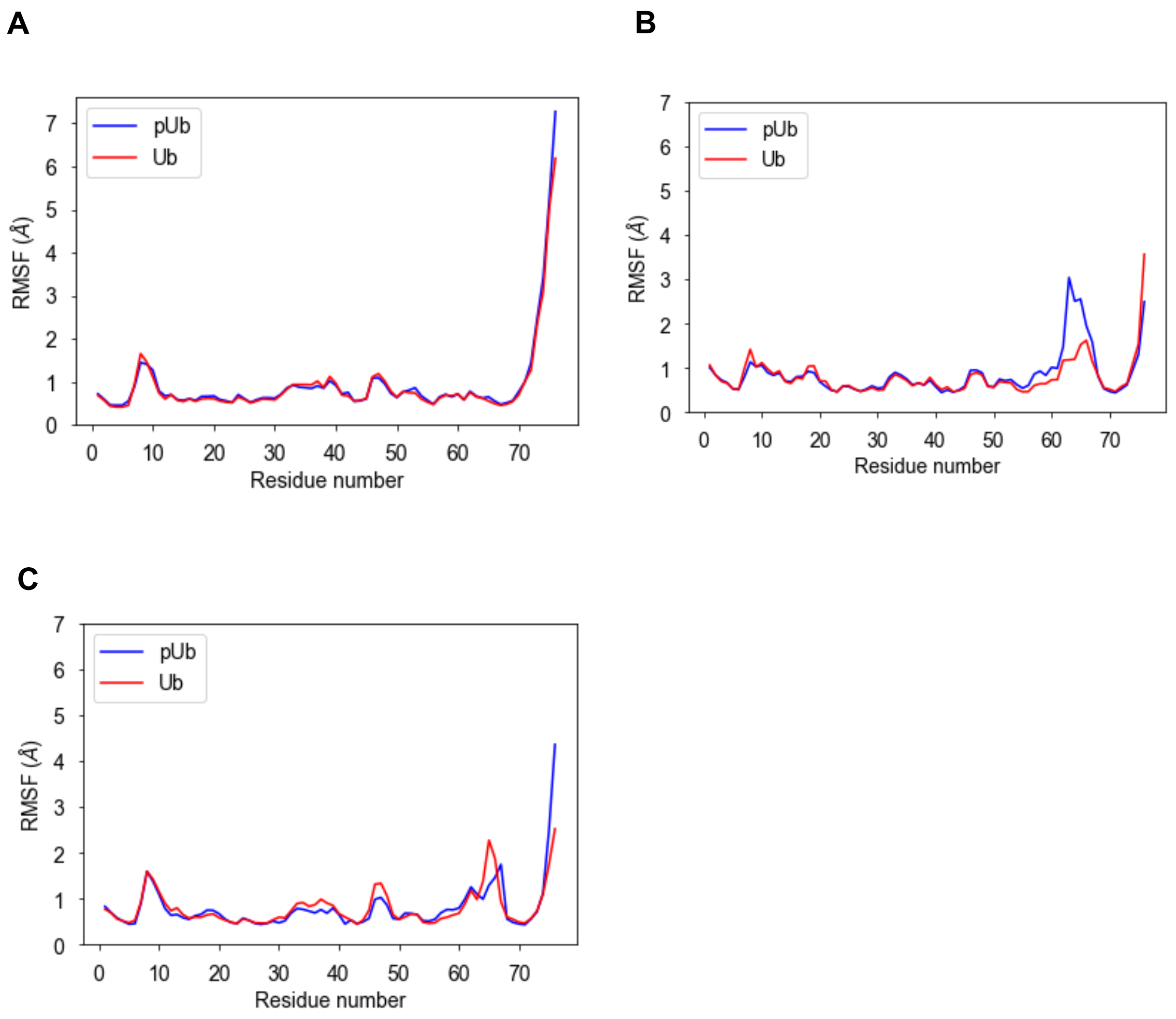


**Fig. S8. RMSF plots by residue, related to Fig. 3.** $C_\alpha$ root-mean-square fluctuations (RMSF) for both pUb (blue) and Ub (red) in the **(A)** Major, **(B)** Bent, and **(C)** CR conformations.

**Table S1 – Correlation data for pUb in the Major conformation**

| $\beta$5 residue | $\beta$1 or $\beta$3 residue | Corr | Corr SEM | #windows |
|---|---|---|---|---|
| resname VAL and resid 70 | resname GLN and resid 40 | 0.4537 | 0.0474 | 4 |
| resname ARG and resid 72 | resname ASP and resid 39 | 0.4278 | 0.0490 | 4 |
| resname ARG and resid 72 | resname GLN and resid 40 | 0.4181 | 0.0511 | 4 |
| resname VAL and resid 70 | resname ARG and resid 42 | 0.4176 | 0.0430 | 4 |
| resname VAL and resid 70 | resname GLN and resid 41 | 0.4128 | 0.0417 | 4 |
| resname LEU and resid 71 | resname GLN and resid 40 | 0.4036 | 0.0496 | 4 |
| resname ARG and resid 72 | resname ARG and resid 42 | 0.3815 | 0.0451 | 4 |
| resname VAL and resid 70 | resname LEU and resid 43 | 0.3780 | 0.0470 | 4 |
| resname LEU and resid 71 | resname ARG and resid 42 | 0.3776 | 0.0472 | 4 |
| resname ARG and resid 72 | resname GLN and resid 41 | 0.3745 | 0.0482 | 4 |
| resname LEU and resid 71 | resname GLN and resid 41 | 0.3622 | 0.0454 | 4 |
| resname HSD and resid 68 | resname ARG and resid 42 | 0.3594 | 0.0396 | 4 |
| resname LYS and resid 63 | resname MET and resid 1 | 0.3558 | 0.0514 | 4 |
| resname GLN and resid 62 | resname MET and resid 1 | 0.3484 | 0.0503 | 4 |
| resname VAL and resid 70 | resname ILE and resid 44 | 0.3462 | 0.0431 | 4 |

| | | | | |
|---|---|---|---|---|
| resname LEU and resid 69 | resname GLN and resid 41 | 0.3379 | 0.0466 | 4 |
| resname GLU and resid 64 | resname PHE and resid 4 | 0.3323 | 0.0615 | 4 |
| resname LEU and resid 69 | resname ARG and resid 42 | 0.3302 | 0.0420 | 4 |
| resname GLU and resid 64 | resname ILE and resid 3 | 0.3265 | 0.0611 | 4 |
| resname VAL and resid 70 | resname LEU and resid 8 | 0.3249 | 0.0543 | 4 |
| resname THR and resid 66 | resname VAL and resid 5 | 0.3202 | 0.0547 | 4 |
| resname GLU and resid 64 | resname GLN and resid 2 | 0.3196 | 0.0606 | 4 |
| resname SER and resid 65 | resname PHE and resid 45 | 0.3169 | 0.0269 | 4 |
| resname THR and resid 66 | resname PHE and resid 4 | 0.3161 | 0.0649 | 4 |
| resname THR and resid 66 | resname PHE and resid 45 | 0.3146 | 0.0376 | 4 |
| resname THR and resid 66 | resname LYS and resid 6 | 0.3117 | 0.0549 | 4 |
| resname LEU and resid 67 | resname PHE and resid 45 | 0.3102 | 0.0357 | 4 |
| resname LEU and resid 67 | resname PHE and resid 4 | 0.3088 | 0.0604 | 4 |
| resname LYS and resid 63 | resname ILE and resid 3 | 0.3087 | 0.0574 | 4 |
| resname HSD and resid 68 | resname LEU and resid 43 | 0.3072 | 0.0442 | 4 |
| resname SER and resid 65 | resname PHE and resid 4 | 0.3066 | 0.0539 | 4 |

| | | | | |
|---|---|---|---|---|
| resname LEU and resid 67 | resname ILE and resid 3 | 0.3043 | 0.0616 | 4 |
| resname LEU and resid 71 | resname LEU and resid 8 | 0.3003 | 0.0480 | 4 |
| resname LYS and resid 63 | resname GLN and resid 2 | 0.2988 | 0.0572 | 4 |
| resname SER and resid 65 | resname ILE and resid 3 | 0.2984 | 0.0546 | 4 |
| resname HSD and resid 68 | resname VAL and resid 5 | 0.2965 | 0.0557 | 4 |
| resname GLN and resid 62 | resname ILE and resid 3 | 0.2958 | 0.0554 | 4 |
| resname LEU and resid 67 | resname VAL and resid 5 | 0.2952 | 0.0556 | 4 |
| resname HSD and resid 68 | resname LYS and resid 6 | 0.2937 | 0.0570 | 4 |
| resname LEU and resid 67 | resname ILE and resid 44 | 0.2932 | 0.0405 | 4 |
| resname LEU and resid 67 | resname LYS and resid 6 | 0.2893 | 0.0534 | 4 |
| resname LEU and resid 69 | resname VAL and resid 5 | 0.2892 | 0.0540 | 4 |
| resname SER and resid 65 | resname GLN and resid 2 | 0.2888 | 0.0537 | 4 |
| resname LEU and resid 67 | resname LEU and resid 43 | 0.2875 | 0.0472 | 4 |
| resname LEU and resid 69 | resname LYS and resid 6 | 0.2873 | 0.0557 | 4 |
| resname LEU and resid 69 | resname LEU and resid 8 | 0.2849 | 0.0516 | 4 |
| resname LEU and resid 69 | resname THR and resid 7 | 0.2818 | 0.0544 | 4 |

| | | | | |
|---|---|---|---|---|
| resname LEU and resid 69 | resname LEU and resid 43 | 0.2733 | 0.0447 | 4 |
| resname HSD and resid 68 | resname PHE and resid 45 | 0.2674 | 0.0428 | 4 |
| resname HSD and resid 68 | resname ILE and resid 44 | 0.2630 | 0.0477 | 4 |
| resname LEU and resid 69 | resname ILE and resid 44 | 0.2488 | 0.0429 | 4 |
| resname THR and resid 66 | resname THR and resid 12 | 0.2080 | 0.0751 | 3 |
| resname LEU and resid 73 | resname GLN and resid 40 | 0.0748 | 0.0748 | 1 |
| resname HSD and resid 68 | resname LEU and resid 8 | 0.0558 | 0.0558 | 1 |
| resname HSD and resid 68 | resname THR and resid 7 | 0.0493 | 0.0493 | 1 |

**Table S2 – Correlation data for pUb in the Bent conformation**

| $\beta$5 residue | $\beta$1 or $\beta$3 residue | Corr | Corr SEM | #windows |
|---|---|---|---|---|
| resname ARG and resid 74 | resname ASP and resid 39 | 0.7503 | 0.0173 | 4 |
| resname ARG and resid 74 | resname GLN and resid 40 | 0.7298 | 0.0181 | 4 |
| resname ARG and resid 74 | resname GLN and resid 41 | 0.6806 | 0.0217 | 4 |
| resname ARG and resid 74 | resname ARG and resid 42 | 0.6661 | 0.0183 | 4 |
| resname GLY and resid 75 | resname GLN and resid 40 | 0.6634 | 0.0126 | 4 |
| resname VAL and resid 70 | resname ARG and resid 42 | 0.5512 | 0.0284 | 4 |

| | | | | |
|---|---|---|---|---|
| resname ARG and resid 72 | resname GLN and resid 40 | 0.5475 | 0.0388 | 4 |
| resname VAL and resid 70 | resname LEU and resid 43 | 0.5279 | 0.0320 | 4 |
| resname ARG and resid 72 | resname ARG and resid 42 | 0.5245 | 0.0309 | 4 |
| resname LEU and resid 73 | resname GLN and resid 40 | 0.5244 | 0.0371 | 4 |
| resname ARG and resid 72 | resname GLN and resid 41 | 0.5211 | 0.0361 | 4 |
| resname LEU and resid 71 | resname ARG and resid 42 | 0.5018 | 0.0299 | 4 |
| resname LEU and resid 71 | resname GLN and resid 41 | 0.5011 | 0.0354 | 4 |
| resname ARG and resid 72 | resname LEU and resid 43 | 0.4936 | 0.0357 | 4 |
| resname LEU and resid 73 | resname ARG and resid 42 | 0.4929 | 0.0328 | 4 |
| resname LEU and resid 71 | resname LEU and resid 43 | 0.4903 | 0.0330 | 4 |
| resname LEU and resid 73 | resname GLN and resid 41 | 0.4869 | 0.0373 | 4 |
| resname LEU and resid 69 | resname LEU and resid 43 | 0.4697 | 0.0291 | 4 |
| resname GLN and resid 62 | resname MET and resid 1 | 0.4615 | 0.0554 | 4 |
| resname VAL and resid 70 | resname ILE and resid 44 | 0.4595 | 0.0396 | 4 |
| resname VAL and resid 70 | resname PHE and resid 45 | 0.4463 | 0.0478 | 4 |
| resname VAL and resid 70 | resname VAL and resid 5 | 0.4435 | 0.0532 | 4 |

| | | | | |
|---|---|---|---|---|
| resname LEU and resid 71 | resname ILE and resid 44 | 0.4374 | 0.0380 | 4 |
| resname ARG and resid 72 | resname ILE and resid 44 | 0.4330 | 0.0419 | 4 |
| resname LEU and resid 71 | resname THR and resid 7 | 0.4288 | 0.0422 | 4 |
| resname VAL and resid 70 | resname LYS and resid 6 | 0.4279 | 0.0482 | 4 |
| resname LEU and resid 71 | resname LEU and resid 8 | 0.4227 | 0.0398 | 4 |
| resname LEU and resid 71 | resname VAL and resid 5 | 0.4214 | 0.0522 | 4 |
| resname ARG and resid 72 | resname LEU and resid 8 | 0.4181 | 0.0491 | 4 |
| resname LEU and resid 73 | resname LEU and resid 8 | 0.4163 | 0.0445 | 4 |
| resname LEU and resid 67 | resname PHE and resid 45 | 0.4157 | 0.0430 | 4 |
| resname LEU and resid 71 | resname LYS and resid 6 | 0.4126 | 0.0453 | 4 |
| resname HSD and resid 68 | resname PHE and resid 45 | 0.4063 | 0.0408 | 4 |
| resname LEU and resid 69 | resname ILE and resid 44 | 0.3975 | 0.0337 | 4 |
| resname HSD and resid 68 | resname VAL and resid 5 | 0.3968 | 0.0598 | 4 |
| resname LEU and resid 69 | resname PHE and resid 45 | 0.3870 | 0.0394 | 4 |
| resname LEU and resid 69 | resname VAL and resid 5 | 0.3833 | 0.0590 | 4 |
| resname HSD and resid 68 | resname LYS and resid 6 | 0.3806 | 0.0560 | 4 |

| | | | | |
|---|---|---|---|---|
| resname LEU and resid 69 | resname LYS and resid 6 | 0.3669 | 0.0561 | 4 |
| resname HSD and resid 68 | resname PHE and resid 4 | 0.3667 | 0.0552 | 4 |
| resname LEU and resid 69 | resname ILE and resid 3 | 0.3575 | 0.0525 | 4 |
| resname LEU and resid 69 | resname PHE and resid 4 | 0.3523 | 0.0552 | 4 |
| resname SER and resid 65 | resname PHE and resid 4 | 0.2234 | 0.1318 | 2 |
| resname LEU and resid 67 | resname PHE and resid 4 | 0.2171 | 0.1290 | 2 |
| resname LYS and resid 63 | resname MET and resid 1 | 0.1122 | 0.1122 | 1 |
| resname GLN and resid 62 | resname ILE and resid 3 | 0.1066 | 0.1066 | 1 |
| resname LYS and resid 63 | resname ILE and resid 3 | 0.1005 | 0.1005 | 1 |
| resname GLU and resid 64 | resname PHE and resid 4 | 0.0985 | 0.0985 | 1 |
| resname LYS and resid 63 | resname GLN and resid 2 | 0.0963 | 0.0963 | 1 |
| resname HSD and resid 68 | resname THR and resid 12 | 0.0843 | 0.0843 | 1 |

**Table S3 – Correlation data for pUb in the CR conformation**

| $\beta$5 residue | $\beta$1 or $\beta$3 residue | Corr | Corr SEM | #windows |
|---|---|---|---|---|
| resname ARG and resid 74 | resname GLN and resid 40 | 0.6812 | 0.0262 | 4 |
| resname ARG and resid 74 | resname GLN and resid 41 | 0.6345 | 0.0304 | 4 |

| | | | | |
|---|---|---|---|---|
| resname ARG and resid 74 | resname ARG and resid 42 | 0.6101 | 0.0297 | 4 |
| resname VAL and resid 70 | resname ARG and resid 42 | 0.4754 | 0.0543 | 4 |
| resname VAL and resid 70 | resname LEU and resid 43 | 0.4535 | 0.0551 | 4 |
| resname ARG and resid 72 | resname GLN and resid 40 | 0.4423 | 0.0499 | 4 |
| resname ARG and resid 72 | resname ARG and resid 42 | 0.4398 | 0.0515 | 4 |
| resname ARG and resid 72 | resname GLN and resid 41 | 0.4357 | 0.0509 | 4 |
| resname LEU and resid 71 | resname ARG and resid 42 | 0.4179 | 0.0670 | 4 |
| resname LEU and resid 71 | resname GLN and resid 41 | 0.4173 | 0.0603 | 4 |
| resname ARG and resid 72 | resname LEU and resid 43 | 0.4147 | 0.0590 | 4 |
| resname LEU and resid 71 | resname LEU and resid 43 | 0.4080 | 0.0556 | 4 |
| resname LEU and resid 73 | resname GLN and resid 40 | 0.4042 | 0.0565 | 4 |
| resname VAL and resid 70 | resname ILE and resid 44 | 0.4017 | 0.0659 | 4 |
| resname LEU and resid 73 | resname ARG and resid 42 | 0.3998 | 0.0613 | 4 |
| resname GLU and resid 64 | resname PHE and resid 45 | 0.3993 | 0.0624 | 4 |
| resname GLN and resid 62 | resname MET and resid 1 | 0.3984 | 0.0471 | 4 |
| resname VAL and resid 70 | resname PHE and resid 45 | 0.3951 | 0.0640 | 4 |

| | | | | |
|---|---|---|---|---|
| resname LEU and resid 73 | resname GLN and resid 41 | 0.3919 | 0.0588 | 4 |
| resname LEU and resid 71 | resname LEU and resid 8 | 0.3765 | 0.0369 | 4 |
| resname LEU and resid 69 | resname LEU and resid 43 | 0.3702 | 0.0566 | 4 |
| resname LEU and resid 71 | resname ILE and resid 44 | 0.3678 | 0.0672 | 4 |
| resname ARG and resid 72 | resname ILE and resid 44 | 0.3668 | 0.0657 | 4 |
| resname THR and resid 66 | resname PHE and resid 45 | 0.3660 | 0.0676 | 4 |
| resname ARG and resid 72 | resname LEU and resid 8 | 0.3629 | 0.0371 | 4 |
| resname LEU and resid 73 | resname LEU and resid 8 | 0.3586 | 0.0420 | 4 |
| resname VAL and resid 70 | resname VAL and resid 5 | 0.3512 | 0.0496 | 4 |
| resname LEU and resid 67 | resname PHE and resid 45 | 0.3443 | 0.0617 | 4 |
| resname VAL and resid 70 | resname LYS and resid 6 | 0.3380 | 0.0420 | 4 |
| resname LEU and resid 71 | resname THR and resid 7 | 0.3374 | 0.0441 | 4 |
| resname LEU and resid 71 | resname VAL and resid 5 | 0.3326 | 0.0481 | 4 |
| resname HSD and resid 68 | resname PHE and resid 45 | 0.3294 | 0.0630 | 4 |
| resname LEU and resid 71 | resname LYS and resid 6 | 0.3227 | 0.0360 | 4 |
| resname LEU and resid 69 | resname ILE and resid 44 | 0.3134 | 0.0570 | 4 |

| | | | | |
|---|---|---|---|---|
| resname LEU and resid 69 | resname PHE and resid 45 | 0.3133 | 0.0597 | 4 |
| resname LEU and resid 69 | resname VAL and resid 5 | 0.2889 | 0.0462 | 4 |
| resname HSD and resid 68 | resname LYS and resid 6 | 0.2874 | 0.0400 | 4 |
| resname LEU and resid 69 | resname LYS and resid 6 | 0.2709 | 0.0366 | 4 |
| resname HSD and resid 68 | resname THR and resid 12 | 0.2702 | 0.0348 | 4 |
| resname LEU and resid 67 | resname PHE and resid 4 | 0.2691 | 0.0400 | 4 |
| resname HSD and resid 68 | resname PHE and resid 4 | 0.2486 | 0.0613 | 4 |
| resname LEU and resid 69 | resname PHE and resid 4 | 0.2287 | 0.0566 | 4 |
| resname LEU and resid 69 | resname ILE and resid 3 | 0.2087 | 0.0578 | 4 |
| resname ARG and resid 74 | resname ASP and resid 39 | 0.5223 | 0.1750 | 3 |
| resname LEU and resid 67 | resname GLN and resid 2 | 0.1559 | 0.0947 | 2 |
| resname HSD and resid 68 | resname VAL and resid 5 | 0.1489 | 0.0904 | 2 |
| resname LEU and resid 67 | resname ILE and resid 3 | 0.1469 | 0.0866 | 2 |
| resname LEU and resid 67 | resname MET and resid 1 | 0.0651 | 0.0651 | 1 |

**Table S4 – Correlation data for Ub in the Major conformation**

| **$\beta$5 residue** | **$\beta$1 or $\beta$3 residue** | **Corr** | **Corr SEM** | **#windows** |
|---|---|---|---|---|
| resname VAL and resid 70 | resname GLN and resid 40 | 0.4489 | 0.0238 | 4 |

| | | | | |
|---|---|---|---|---|
| resname ARG and resid 72 | resname GLN and resid 40 | 0.4154 | 0.0246 | 4 |
| resname LEU and resid 71 | resname GLN and resid 40 | 0.3979 | 0.0332 | 4 |
| resname VAL and resid 70 | resname GLN and resid 41 | 0.3966 | 0.0188 | 4 |
| resname VAL and resid 70 | resname ARG and resid 42 | 0.3799 | 0.0210 | 4 |
| resname ARG and resid 72 | resname GLN and resid 41 | 0.3556 | 0.0186 | 4 |
| resname VAL and resid 70 | resname LEU and resid 43 | 0.3492 | 0.0115 | 4 |
| resname LEU and resid 71 | resname GLN and resid 41 | 0.3393 | 0.0256 | 4 |
| resname ARG and resid 72 | resname ARG and resid 42 | 0.3352 | 0.0208 | 4 |
| resname LEU and resid 71 | resname ARG and resid 42 | 0.3337 | 0.0269 | 4 |
| resname HSD and resid 68 | resname ARG and resid 42 | 0.3336 | 0.0088 | 4 |
| resname LYS and resid 63 | resname MET and resid 1 | 0.3260 | 0.0126 | 4 |
| resname VAL and resid 70 | resname LEU and resid 8 | 0.3124 | 0.0099 | 4 |
| resname LEU and resid 69 | resname GLN and resid 41 | 0.3029 | 0.0082 | 4 |
| resname THR and resid 66 | resname VAL and resid 5 | 0.2931 | 0.0106 | 4 |
| resname GLU and resid 64 | resname PHE and resid 4 | 0.2926 | 0.0311 | 4 |
| resname LEU and resid 69 | resname ARG and resid 42 | 0.2923 | 0.0048 | 4 |

| | | | | |
|---|---|---|---|---|
| resname VAL and resid 70 | resname ILE and resid 44 | 0.2896 | 0.0034 | 4 |
| resname LYS and resid 63 | resname ILE and resid 3 | 0.2887 | 0.0213 | 4 |
| resname LEU and resid 71 | resname LEU and resid 8 | 0.2878 | 0.0138 | 4 |
| resname GLU and resid 64 | resname GLN and resid 2 | 0.2862 | 0.0297 | 4 |
| resname GLU and resid 64 | resname ILE and resid 3 | 0.2861 | 0.0308 | 4 |
| resname LEU and resid 67 | resname LEU and resid 43 | 0.2853 | 0.0130 | 4 |
| resname LYS and resid 63 | resname GLN and resid 2 | 0.2833 | 0.0233 | 4 |
| resname LEU and resid 67 | resname VAL and resid 5 | 0.2820 | 0.0110 | 4 |
| resname SER and resid 65 | resname GLN and resid 2 | 0.2810 | 0.0292 | 4 |
| resname HSD and resid 68 | resname LEU and resid 43 | 0.2789 | 0.0085 | 4 |
| resname THR and resid 66 | resname LYS and resid 6 | 0.2731 | 0.0106 | 4 |
| resname LEU and resid 67 | resname LYS and resid 6 | 0.2701 | 0.0115 | 4 |
| resname SER and resid 65 | resname PHE and resid 4 | 0.2695 | 0.0353 | 4 |
| resname THR and resid 66 | resname PHE and resid 4 | 0.2617 | 0.0337 | 4 |
| resname SER and resid 65 | resname ILE and resid 3 | 0.2605 | 0.0352 | 4 |
| resname HSD and resid 68 | resname VAL and resid 5 | 0.2560 | 0.0257 | 4 |

| | | | | |
|---|---|---|---|---|
| resname SER and resid 65 | resname PHE and resid 45 | 0.2550 | 0.0121 | 4 |
| resname HSD and resid 68 | resname LYS and resid 6 | 0.2542 | 0.0237 | 4 |
| resname LEU and resid 67 | resname PHE and resid 4 | 0.2538 | 0.0334 | 4 |
| resname LEU and resid 67 | resname ILE and resid 3 | 0.2529 | 0.0363 | 4 |
| resname LEU and resid 69 | resname LEU and resid 43 | 0.2512 | 0.0133 | 4 |
| resname THR and resid 66 | resname THR and resid 12 | 0.2443 | 0.0148 | 4 |
| resname LEU and resid 69 | resname THR and resid 7 | 0.2402 | 0.0256 | 4 |
| resname LEU and resid 69 | resname VAL and resid 5 | 0.2402 | 0.0245 | 4 |
| resname LEU and resid 67 | resname ILE and resid 44 | 0.2343 | 0.0167 | 4 |
| resname LEU and resid 69 | resname LYS and resid 6 | 0.2341 | 0.0235 | 4 |
| resname LEU and resid 69 | resname LEU and resid 8 | 0.2309 | 0.0262 | 4 |
| resname LEU and resid 67 | resname PHE and resid 45 | 0.2301 | 0.0168 | 4 |
| resname THR and resid 66 | resname PHE and resid 45 | 0.2293 | 0.0103 | 4 |
| resname HSD and resid 68 | resname ILE and resid 44 | 0.2090 | 0.0112 | 4 |
| resname HSD and resid 68 | resname PHE and resid 45 | 0.2026 | 0.0067 | 4 |
| resname LEU and resid 69 | resname ILE and resid 44 | 0.1918 | 0.0070 | 4 |

| | | | | |
|---|---|---|---|---|
| resname ARG and resid 72 | resname ASP and resid 39 | 0.3190 | 0.1080 | 3 |
| resname ARG and resid 74 | resname GLN and resid 40 | 0.1436 | 0.1436 | 1 |

**Table S5 – Correlation data for Ub in the Bent conformation**

| $\beta$5 residue | $\beta$1 or $\beta$3 residue | Corr | Corr SEM | #windows |
|---|---|---|---|---|
| resname ARG and resid 74 | resname ASP and resid 39 | 0.7398 | 0.0164 | 4 |
| resname ARG and resid 74 | resname GLN and resid 40 | 0.7199 | 0.0236 | 4 |
| resname ARG and resid 74 | resname GLN and resid 41 | 0.6628 | 0.0209 | 4 |
| resname GLY and resid 75 | resname GLN and resid 40 | 0.6596 | 0.0298 | 4 |
| resname ARG and resid 74 | resname ARG and resid 42 | 0.6418 | 0.0175 | 4 |
| resname VAL and resid 70 | resname ARG and resid 42 | 0.5615 | 0.0289 | 4 |
| resname VAL and resid 70 | resname LEU and resid 43 | 0.5332 | 0.0285 | 4 |
| resname ARG and resid 72 | resname ARG and resid 42 | 0.5324 | 0.0330 | 4 |
| resname ARG and resid 72 | resname GLN and resid 41 | 0.5278 | 0.0223 | 4 |
| resname LEU and resid 71 | resname ARG and resid 42 | 0.5244 | 0.0237 | 4 |
| resname LEU and resid 73 | resname GLN and resid 40 | 0.5229 | 0.0186 | 4 |
| resname LEU and resid 71 | resname GLN and resid 41 | 0.5109 | 0.0115 | 4 |

| | | | | |
|---|---|---|---|---|
| resname LEU and resid 73 | resname ARG and resid 42 | 0.5030 | 0.0305 | 4 |
| resname ARG and resid 72 | resname LEU and resid 43 | 0.5024 | 0.0354 | 4 |
| resname LEU and resid 71 | resname LEU and resid 43 | 0.4993 | 0.0241 | 4 |
| resname LEU and resid 73 | resname GLN and resid 41 | 0.4907 | 0.0168 | 4 |
| resname LEU and resid 69 | resname LEU and resid 43 | 0.4846 | 0.0304 | 4 |
| resname VAL and resid 70 | resname PHE and resid 45 | 0.4709 | 0.0400 | 4 |
| resname VAL and resid 70 | resname ILE and resid 44 | 0.4687 | 0.0382 | 4 |
| resname ARG and resid 72 | resname ILE and resid 44 | 0.4428 | 0.0411 | 4 |
| resname LEU and resid 71 | resname ILE and resid 44 | 0.4375 | 0.0366 | 4 |
| resname HSD and resid 68 | resname PHE and resid 45 | 0.4268 | 0.0411 | 4 |
| resname LEU and resid 67 | resname PHE and resid 45 | 0.4247 | 0.0416 | 4 |
| resname ARG and resid 72 | resname LEU and resid 8 | 0.4110 | 0.0542 | 4 |
| resname LEU and resid 71 | resname LEU and resid 8 | 0.4087 | 0.0518 | 4 |
| resname LEU and resid 69 | resname ILE and resid 44 | 0.4084 | 0.0373 | 4 |
| resname LEU and resid 69 | resname PHE and resid 45 | 0.4083 | 0.0406 | 4 |
| resname LEU and resid 73 | resname LEU and resid 8 | 0.4044 | 0.0543 | 4 |

| | | | | |
|---|---|---|---|---|
| resname VAL and resid 70 | resname VAL and resid 5 | 0.3938 | 0.0420 | 4 |
| resname LEU and resid 71 | resname THR and resid 7 | 0.3921 | 0.0409 | 4 |
| resname HSD and resid 68 | resname VAL and resid 5 | 0.3879 | 0.0323 | 4 |
| resname LEU and resid 69 | resname VAL and resid 5 | 0.3781 | 0.0270 | 4 |
| resname LEU and resid 71 | resname VAL and resid 5 | 0.3671 | 0.0376 | 4 |
| resname VAL and resid 70 | resname LYS and resid 6 | 0.3671 | 0.0371 | 4 |
| resname HSD and resid 68 | resname LYS and resid 6 | 0.3544 | 0.0268 | 4 |
| resname LEU and resid 69 | resname LYS and resid 6 | 0.3445 | 0.0270 | 4 |
| resname LEU and resid 71 | resname LYS and resid 6 | 0.3376 | 0.0385 | 4 |
| resname HSD and resid 68 | resname PHE and resid 4 | 0.3178 | 0.0341 | 4 |
| resname LEU and resid 69 | resname PHE and resid 4 | 0.3067 | 0.0325 | 4 |
| resname LEU and resid 69 | resname ILE and resid 3 | 0.2980 | 0.0375 | 4 |
| resname ARG and resid 72 | resname GLN and resid 40 | 0.2603 | 0.1503 | 2 |
| resname HSD and resid 68 | resname THR and resid 12 | 0.1060 | 0.1060 | 1 |

**Table S6 – Correlation data for Ub in the CR conformation**

| **$\beta$5 residue** | **$\beta$1 or $\beta$3 residue** | **Corr** | **Corr SEM** | **#windows** |
|---|---|---|---|---|

| | | | | |
|---|---|---|---|---|
| resname ARG and resid 74 | resname ASP and resid 39 | 0.6991 | 0.0067 | 4 |
| resname ARG and resid 74 | resname GLN and resid 40 | 0.6652 | 0.0024 | 4 |
| resname ARG and resid 74 | resname GLN and resid 41 | 0.6074 | 0.0115 | 4 |
| resname ARG and resid 74 | resname ARG and resid 42 | 0.5808 | 0.0084 | 4 |
| resname VAL and resid 70 | resname ARG and resid 42 | 0.4751 | 0.0310 | 4 |
| resname VAL and resid 70 | resname LEU and resid 43 | 0.4598 | 0.0352 | 4 |
| resname LEU and resid 71 | resname ARG and resid 42 | 0.4355 | 0.0317 | 4 |
| resname LEU and resid 71 | resname LEU and resid 43 | 0.4348 | 0.0333 | 4 |
| resname ARG and resid 72 | resname ARG and resid 42 | 0.4231 | 0.0304 | 4 |
| resname LEU and resid 71 | resname GLN and resid 41 | 0.4148 | 0.0135 | 4 |
| resname ARG and resid 72 | resname GLN and resid 41 | 0.4137 | 0.0117 | 4 |
| resname LEU and resid 73 | resname GLN and resid 40 | 0.4096 | 0.0122 | 4 |
| resname ARG and resid 72 | resname LEU and resid 43 | 0.4068 | 0.0391 | 4 |
| resname LEU and resid 73 | resname ARG and resid 42 | 0.3998 | 0.0377 | 4 |
| resname VAL and resid 70 | resname PHE and resid 45 | 0.3948 | 0.0383 | 4 |
| resname VAL and resid 70 | resname ILE and resid 44 | 0.3904 | 0.0408 | 4 |

| | | | | |
|---|---|---|---|---|
| resname LEU and resid 73 | resname GLN and resid 41 | 0.3854 | 0.0124 | 4 |
| resname LEU and resid 71 | resname ILE and resid 44 | 0.3818 | 0.0412 | 4 |
| resname LEU and resid 67 | resname PHE and resid 45 | 0.3597 | 0.0439 | 4 |
| resname LEU and resid 69 | resname LEU and resid 43 | 0.3591 | 0.0263 | 4 |
| resname ARG and resid 72 | resname ILE and resid 44 | 0.3537 | 0.0416 | 4 |
| resname LEU and resid 71 | resname LEU and resid 8 | 0.3423 | 0.0287 | 4 |
| resname LEU and resid 71 | resname THR and resid 7 | 0.3171 | 0.0311 | 4 |
| resname VAL and resid 70 | resname VAL and resid 5 | 0.3159 | 0.0313 | 4 |
| resname HSD and resid 68 | resname PHE and resid 45 | 0.3086 | 0.0285 | 4 |
| resname VAL and resid 70 | resname LYS and resid 6 | 0.3061 | 0.0384 | 4 |
| resname ARG and resid 72 | resname LEU and resid 8 | 0.3041 | 0.0269 | 4 |
| resname LEU and resid 71 | resname VAL and resid 5 | 0.2991 | 0.0352 | 4 |
| resname LEU and resid 71 | resname LYS and resid 6 | 0.2990 | 0.0415 | 4 |
| resname LEU and resid 69 | resname ILE and resid 44 | 0.2947 | 0.0244 | 4 |
| resname HSD and resid 68 | resname LYS and resid 6 | 0.2898 | 0.0311 | 4 |
| resname LEU and resid 69 | resname PHE and resid 45 | 0.2825 | 0.0299 | 4 |

| | | | | |
|---|---|---|---|---|
| resname LEU and resid 69 | resname LYS and resid 6 | 0.2736 | 0.0325 | 4 |
| resname LEU and resid 67 | resname ILE and resid 3 | 0.2719 | 0.0259 | 4 |
| resname LEU and resid 67 | resname PHE and resid 4 | 0.2708 | 0.0239 | 4 |
| resname LEU and resid 69 | resname VAL and resid 5 | 0.2696 | 0.0283 | 4 |
| resname LEU and resid 69 | resname ILE and resid 3 | 0.2109 | 0.0304 | 4 |
| resname HSD and resid 68 | resname PHE and resid 4 | 0.2010 | 0.0248 | 4 |
| resname LEU and resid 69 | resname PHE and resid 4 | 0.1922 | 0.0260 | 4 |
| resname ARG and resid 72 | resname GLN and resid 40 | 0.3250 | 0.1085 | 3 |
| resname THR and resid 66 | resname PHE and resid 45 | 0.2694 | 0.1016 | 3 |
| resname LEU and resid 73 | resname LEU and resid 8 | 0.2365 | 0.0859 | 3 |
| resname HSD and resid 68 | resname THR and resid 12 | 0.2231 | 0.0830 | 3 |
| resname LEU and resid 67 | resname MET and resid 1 | 0.2055 | 0.0720 | 3 |
| resname LEU and resid 67 | resname GLN and resid 2 | 0.1995 | 0.0706 | 3 |
| resname GLY and resid 75 | resname ARG and resid 42 | 0.1270 | 0.1270 | 1 |
| resname HSD and resid 68 | resname VAL and resid 5 | 0.0905 | 0.0905 | 1 |
| resname THR and resid 66 | resname PHE and resid 4 | 0.0734 | 0.0734 | 1 |
